# A disease-associated PPP2R3C-MAP3K1 phospho-regulatory module controls centrosome function

**DOI:** 10.1101/2024.04.02.587836

**Authors:** Anil Kumar Ganga, Lauren K. Sweeney, Armando Rubio Ramos, Cassandra S. Bishop, Virginie Hamel, Paul Guichard, David K. Breslow

## Abstract

Centrosomes have critical roles in microtubule organization and in cell signaling.^1–8^ However, the mechanisms that regulate centrosome function are not fully defined, and thus how defects in centrosomal regulation contribute to disease is incompletely understood. From functional genomic analyses, we find here that PPP2R3C, a PP2A phosphatase subunit, is a distal centriole protein and functional partner of centriolar proteins CEP350 and FOP. We further show that a key function of PPP2R3C is to counteract the kinase activity of MAP3K1. In support of this model, *MAP3K1* knockout suppresses growth defects caused by *PPP2R3C* inactivation, and MAP3K1 and PPP2R3C have opposing effects on basal and microtubule stress-induced JNK signaling. Illustrating the importance of balanced MAP3K1 and PPP2R3C activities, acute overexpression of MAP3K1 severely inhibits centrosome function and triggers rapid centriole disintegration. Additionally, inactivating *PPP2R3C* mutations and activating *MAP3K1* mutations both cause congenital syndromes characterized by gonadal dysgenesis.^9–15^ As a syndromic *PPP2R3C* variant is defective in centriolar localization and binding to centriolar protein FOP, we propose that imbalanced activity of this centrosomal kinase-phosphatase pair is the shared cause of these disorders. Thus, our findings reveal a new centrosomal phospho-regulatory module, shed light on disorders of gonadal development, and illustrate the power of systems genetics to identify previously unrecognized gene functions.

## Main

Centrosomes have critical cellular roles in patterning the microtubule cytoskeleton, templating primary cilia, regulating cell polarity, and organizing cell signaling.^1–8^ Consisting of a pair of centrioles and the surrounding pericentriolar material (PCM), centrosomes contain over 150 identified protein components that underlie their structural and functional complexity.^16,17^ Additionally, centrosomes are subject to intricate regulatory mechanisms that govern their duplication, maturation, maintenance, and diverse cellular functions.^2–4,6,18^

As for other cellular processes, kinases and phosphatases have key roles in regulation of centrosome function. For example, Plk4, Plk1, Nek2, and AurA are critical for centriole duplication, maturation and remodeling of centriolar appendages, and centrosomal microtubule nucleation.^2,3,18–21^ Conversely, reduced activity of Polo kinases can promote developmental centrosome inactivation and centriole loss.^22^ In parallel to these kinases, phosphatases such as calcineurin, Cdc14a, and PP2A (Protein Phosphatase 2A) also control key centrosomal activities.^23–26^ PP2A phosphatases represent a series of holoenzyme complexes in which catalytic subunits associate with one of many regulatory subunits that control PP2A activity, localization, and substrate specificity.^27–29^ Among PP2A holoenzymes, some localize to centrosomes and regulate centrosome function. In *C. elegans* and *D. melanogaster*, PP2A acts with B55 regulatory to ensure centriole duplication and to control mitotic PCM remodeling.^30–34^ In *C. elegans*, RSA-1 (homolog of human PPP2R3A/B/C) regulates mitotic spindle assembly and centrosomal microtubule nucleation, in part by restraining the activity of depolymerizing kinesins.^35^ In mammals, the B56α regulatory subunit (PPP2R5A) localizes to centrosomes,^36^ and the endogenously tagged PPP2CA catalytic was observed at centrosomes in a recent large-scale study.^37^ However, the functions of specific PP2A enzyme complexes at centrosomes in mammalian cells remain poorly understood.

The broader importance of understanding how centrosomal functions are regulated is highlighted by the many disease conditions that result from altered centrosome function.^38^ For example, inherited defects in centriolar proteins impair brain development and cause microcephaly^39,40^, while altered structure or number of centrioles is a common feature of cancer cells and can promote tumorigenesis.^41,42^ Additionally, mutations in genes encoding some centriolar proteins impair ciliary function and cause ciliopathies.^43^ Understanding these pathologies has helped to reveal fundamental features of centrosome biology, to advance disease understanding, and to identify therapeutic strategies that can mitigate or exploit centrosomal alterations.^44,45^ Given the myriad functions of centrioles, a question that remains to be fully answered is whether additional disorders that have been defined clinically can also be attributed mechanistically to centrosome dysfunction.

### Ppp2r3c is a centriolar protein associated with FOP and CEP350

We initially identified the poorly characterized PP2A regulatory subunit gene *Ppp2r3c* as a hit in a genome-wide CRISPR knockout (KO) screen for regulators of signaling in the Hedgehog (Hh) pathway, a process that is dependent on primary cilia in vertebrates (Fig. 1A).^46^ While many hits from this screen are specifically required for ciliary function, several encode centriolar proteins that reflect the essential role of the mother centriole in cilia formation and function (Fig. 1A). To investigate PPP2R3C function, we made use of the DepMap functional genomic resource, in which genome-wide CRISPR KO screens have been conducted in >1000 human cell lines.^47^ For *PPP2R3C*, gene KO had highly variable effects on fitness across cell lines, ranging from minimal growth phenotypes to strong defects (Fig. S1A). These results differ from ciliary genes such as *IFT88*, which have minimal growth phenotypes in this dataset. We therefore asked if any other genes exhibit a similar pattern of variation in growth phenotype across cell lines, as such co-essentiality relationships provide strong predictions of shared function.^46,48–51^ Among 16,708 genes analyzed, growth phenotypes for *FOP* (also known as *FGFR1OP* and *CEP43*) and *CEP350* were most highly correlated to those of *PPP2R3C* (Fig. 1B-C). As FOP and CEP350 are both established centriolar proteins and physically interact with each other,^52^ we reasoned that PPP2R3C may function together with these proteins at centrioles.

**Figure 1.**
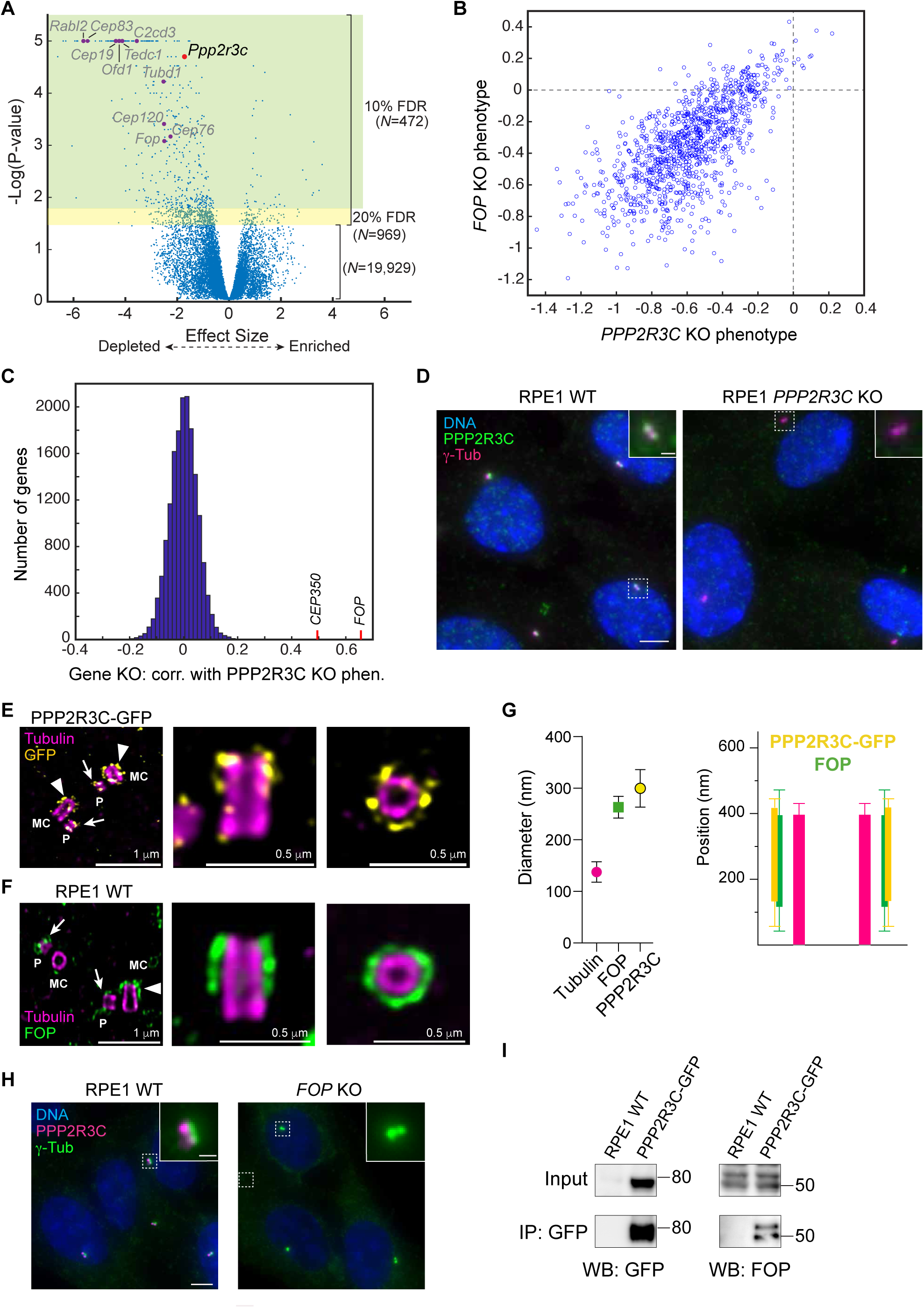
PPP2R3C is a centriolar protein linked to FOP and CEP350. **A)** Hit genes from a genome-wide CRISPR screen for regulators of cilium-dependent Hedgehog signaling are shown, including *Ppp2r3c* as well as several genes that encode centriolar proteins. **B)** CRISPR knockout growth phenotypes across >1000 cancer cell lines in the DepMap dataset are shown for *PPP2R3C* and *FOP* genes, revealing a strong correlation. **C**) Histogram showing correlation coefficients calculated between PPP2R3C’s fitness profile across DepMap CRISPR knockout screens and each of >17,000 other genes in the DepMap dataset. *CEP350* and *FOP* are highlighted as the two genes with fitness profiles most highly correlated to that of *PPP2R3C*. **D)** Immunofluorescence microscopy analysis of PPP2R3C and centrosomes (marked by ψ-tubulin; ψ-tub) in wildtype RPE1 cells and *PPP2R3C* knockout (KO) cells. One of *N* = 3 representative experiments. Scale bar: 5 μm; inset: 1 μm. **E-F)** Representative ultra-structured expansion microscopy (U-ExM) images of RPE1 PPP2R3C-GFP cells stained for GFP (yellow) and tubulin (magenta) (**E**) or wildtype RPE1 cells stained for tubulin (magenta) and FOP (green) (**F**). Labels indicate the mother centrioles (MC; arrowheads) and the procentrioles (P; arrows). Scale bars: 1 and 0.5 μm (pre-expansion). **G)** Relative positions of the tubulin wall, PPP2R3C-GFP (yellow) and FOP (green) at the centriole are shown. Quantifications are based on n=38 centrioles from 3 independent experiments (PPP2R3C-GFP) and n=37 centrioles from 3 independent experiments (FOP). Error bars indicate standard deviation. **H)** Immunofluorescence microscopy analysis of PPP2R3C and centrosomes (marked by ψ-tubulin; ψ-tub) in wildtype RPE1 cells and *FOP* KO cells. One of *N* = 3 representative experiments. Scale bar: 5 μm; inset: 1 μm. **I)** Proteins captured via anti-GFP immunoprecipitation from wildtype RPE1 cells and from RPE1 cells stably expressing PPP2R3C-GFP were detected by immunoblotting, as indicated. One of *N* = 3 representative experiments.

Consistent with this model, we observed prominent localization of PPP2R3C to mother and daughter centrioles in RPE1 cells (Fig. 1D). This localization was observed via an antibody that specifically detects endogenous PPP2R3C, as staining is lost in *PPP2R3C* KO cells. To further characterize PPP2R3C localization at centrioles, we used ultra-structure expansion microscopy (U-ExM), a method that achieves ∼4-fold expansion of biological specimens by embedding them in a swellable hydrogel, thereby revealing features that cannot be resolved by conventional light microscopy.^53^ Using U-ExM, we observed a punctate localization pattern around both centrioles for endogenous PPP2R3C, possibly reflecting compromised antibody staining after gelation and denaturation (Fig. S1B). As we observed more robust labeling via stable expression of a PPP2R3C-GFP transgene (which we validated as functional; see below), we used PPP2R3C-GFP and anti-GFP staining for further analysis. U-ExM of PPP2R3C-GFP revealed a cylindrical distribution at the distal end of centrioles with a diameter of ∼300 nm and length of ∼250 nm (Fig. 1E, G). PPP2R3C-GFP was also observed at the distal end of pro-centrioles, indicating that it is recruited soon after new centrioles form. Notably, this localization pattern precisely matches the pattern observed for FOP and is highly similar to the reported localization of CEP350,^54^ supporting the predicted close functional relationship between these proteins (Fig. 1F-G). The observed centriolar localization is further supported by the identification of PPP2R3C in two mass spectrometry efforts to define a centrosomal proteome.^17,55^

To address the functional hierarchy underlying the shared localization of PPP2R3C and FOP, we analyzed PPP2R3C localization in *FOP* KO cells and observed complete loss of centriolar staining (Fig. 1H). Conversely, FOP localization at centrioles was preserved in *PPP2R3C* KO cells (but absent in *FOP* KO cells) (Fig. S1C-E). In addition to confirming the specificity of FOP staining, these findings indicate that FOP localizes to centrioles independently of PPP2R3C but is needed to recruit PPP2R3C to centrioles. Lastly, we immunoprecipitated PPP2R3C-GFP from RPE1 cells and detected robust co-precipitation of FOP with PPP2R3C-GFP (Fig. 1I). Together, these functional genomic, microscopy, and biochemical data identify PPP2R3C as a centriolar protein associated with FOP and CEP350.

### MAP3K1 is a novel centriolar kinase linked to PPP2R3C

Although variation in the *PPP2R3C* mutant phenotype helped us to predict a close functional relationship with *FOP* and *CEP350*, a key question is why knockout of these genes is tolerated in some cell lines (e.g. RPE1) but not in others. To gain insight into this variability, we asked if cell line-specific differences in the expression of any gene might be predictive of the effect of *PPP2R3C* inactivation. Strikingly, among 19,220 genes with available expression data, *MAP3K1* stood out with its expression levels strongly correlated with the severity of the *PPP2R3C* KO growth defect (Fig. 2A-B). This trend was observed across cancer types, with some subtypes such as neuroblastoma showing particularly high MAP3K1 expression and strong dependence on PPP2R3C (Fig. S2A). *MAP3K1* (also known as *MEKK1*) encodes a MEKK family kinase that activates JNK signaling under stress conditions, including microtubule perturbations.^56,57^ Notably, this relationship between *PPP2R3C* and *MAP3K1* across cancer cell lines is consistent with the antagonistic enzymatic activities of their gene products and suggests that centrosomal PPP2R3C-PP2A phosphatase opposes MAP3K1-mediated phosphorylation.

**Figure 2.**
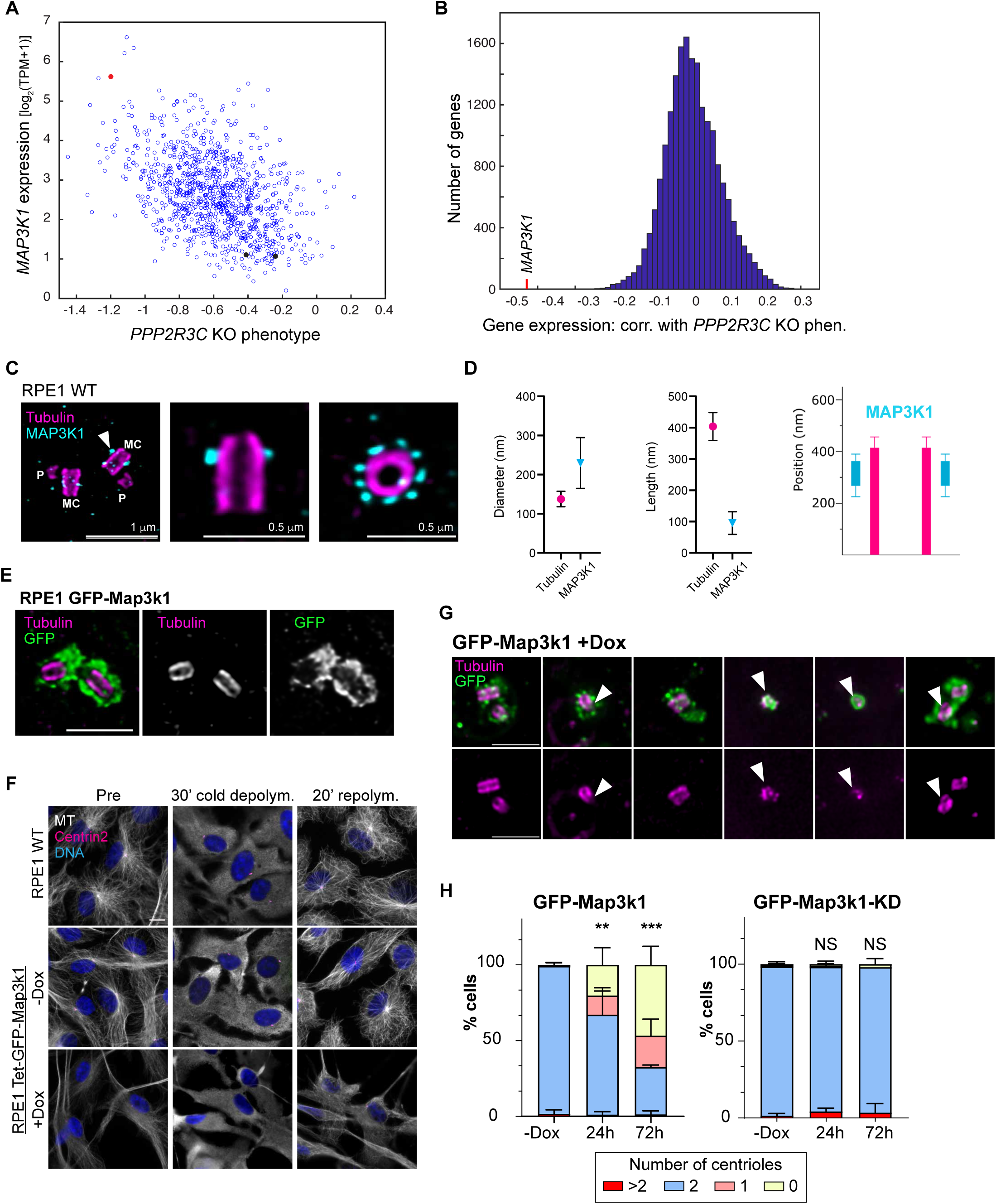
MAP3K1 is a centrosomal kinase that negatively regulates centrosome function. **A)** Relationship between expression levels of *MAP3K1* and fitness phenotype of *PPP2R3C* KO across >1000 cancer lines in the DepMap dataset. Red dot indicates SKNBE2 neuroblastoma cells and black dots indicate two diploid RPE1 clones. **B)** Histogram showing correlation coefficients between the expression level of each gene in the genome across DepMap cancer cell lines and the *PPP2R3C* KO fitness phenotype in these cells lines. MAP3K1 expression is negatively correlated (higher *MAP3K1* expression is associated with greater reduction in fitness upon *PPP2R3C* KO. **C)** Representative U-ExM image of RPE1 cells stained for tubulin (magenta) and MAP3K1 (cyan), with nine-fold symmetric MAP3K1 localization revealed in top view (right). MC: mother centriole, arrowhead; P: pro-centriole. Scale bars: 1 and 0.5 µm (pre-expansion). **D)** Relative positions of the centriole tubulin wall and MAP3K1 were quantified for n=19 centrioles from 2 independent experiments. Error bars indicate standard deviation. **E)** Representative U-ExM images of RPE1 cells expressing GFP-Map3k1 stained for tubulin and GFP. Scale bar: 1 µm (pre-expansion). **F)** Immunofluorescence staining shows microtubule organization before, during, and 20 min after cold-induced depolymerization of microtubules. Images show wildtype RPE1 cells or RPE1 cells with Dox-inducible expression of GFP-Map3k1. Scale bar: 10 μm. See also Fig. S2C. **G)** Representative images of deformed and/or disintegrating centrioles observed in cells over-expressing GFP-Map3k1 for 24-36 hrs. Arrowheads denote centrioles with structural defects. Scale bar: 1 µm (pre-expansion). **H)** Quantification of centriole number per cell in cells over-expressing GFP-Map3k1 (left) or kinase-dead (KD) Map3k1-GFP for the indicated duration (n > 200 cells per cell line from N = 3 independent experiments. Error bars indicate standard deviation. Asterisks denote significant differences in mean (**, *P* = 0.0001; ***, *P* < 0.0001; NS, not significant).

To test this model for PPP2R3C and MAP3K1 function, we pursued a strategy to experimentally increase *MAP3K1* levels in RPE1 cells, which have low intrinsic *MAP3K1* expression in the DepMap dataset (Fig. 2A). We stably transduced wildtype RPE1 cells with a tetracycline-inducible GFP-Map3k1 transgene and observed robust expression upon doxycycline addition. Notably, GFP-Map3k1 exhibited prominent localization at centrioles (Fig. S2B). We therefore used U-ExM to visualize endogenous Map3k1 in RPE1 cells and observed a 9-fold symmetric pattern on the mother centriole that closely resembles that of sub-distal appendage proteins (Fig. 2C-D). We similarly analyzed the localization of over-expressed GFP-Map3k1 by U-ExM and observed a broadened localization around both the mother and daughter centrioles in a pattern reminiscent of the pericentriolar material (PCM) (Fig. 2E).

Having established Map3k1 as a centrosomal kinase, we next assessed the functional impact of GFP-Map3k1 overexpression on centrosomal activity. We first examined centrosomal microtubule nucleation after depolymerization of microtubules by cold shock. Upon Map3k1 overexpression, cells exhibited a severe defect in microtubule nucleation from centrosomes that was not observed in uninduced cells or in cells expressing a kinase-dead construct (Tet-GFP-Map3k1-KD) (Figs. 2F and S2C). Further indicating defective centrosome function, there was also a significant defect in cilium formation in serum-starved cells over-expressing GFP-Map3k1 (Fig. S2D). To investigate the potential basis for these defects, we examined centriole ultra-structure in GFP-Map3k1-expressing cells. Strikingly, we observed pronounced defects in centriole structure, with many centrioles exhibiting a loss of structural integrity and fragmenting (Fig. 2G). Additionally, in many cells centrioles were completely absent, as assessed by U-ExM staining for tubulin (Fig. 2H). Progressive centriole loss was observed over several days but could be observed within 24 hr of GFP-Map3k1 induction, indicating a rapid destabilization of centriole structure. As above, these defects were not observed in uninduced cells or upon expression of Map3k1-KD, even though kinase-dead GFP-Map3k1 still surrounds the centrosomes (Figs. 2H and S2E). Thus, the observed centriolar defects require Map3k1 kinase activity. Lastly, consistent with the severe centrosomal defects observed, we also found that overexpression of Map3k1 but not Map3k1-KD caused cell cycle arrest in G0/G1 within 24 hr of induction (Fig. S2F). Thus, imbalanced activity of MAP3K1 and PPP2R3C causes centrosome dysfunction and strong impairment of cell proliferation.

### Functionally opposing roles for PPP2R3C and MAP3K1

In parallel to increasing *MAP3K1* levels in cells with low intrinsic expression, we pursued a complementary strategy of assessing PPP2R3C function in cells that have high *MAP3K1* expression. To this end, we first examined the effect of *PPP2R3C* knockout on cell proliferation in the SKNBE2 neuroblastoma cell line, as DepMap data indicate high *MAP3K1* expression in these cells (Fig. 3A). We therefore established SKNBE2 cells that express Cas9 in a tetracycline-inducible manner and transduced these cells with an sgRNA cassette that expresses two sgRNAs targeting *PPP2R3C* as well as mCherry to mark transduced cells. Following induction of Cas9, we used serial flow cytometry analysis to assess the relative growth of transduced mCherry-positive cells (Fig. 3A). Notably, we observed strong depletion of mCherry-positive cells over time, while mCherry-positive cells were maintained at stable levels in uninduced cells not expressing Cas9 (Fig. 3B). Thus, *PPP2R3C* KO strongly impairs cell proliferation in SKNBE2 cells.

**Figure 3.**
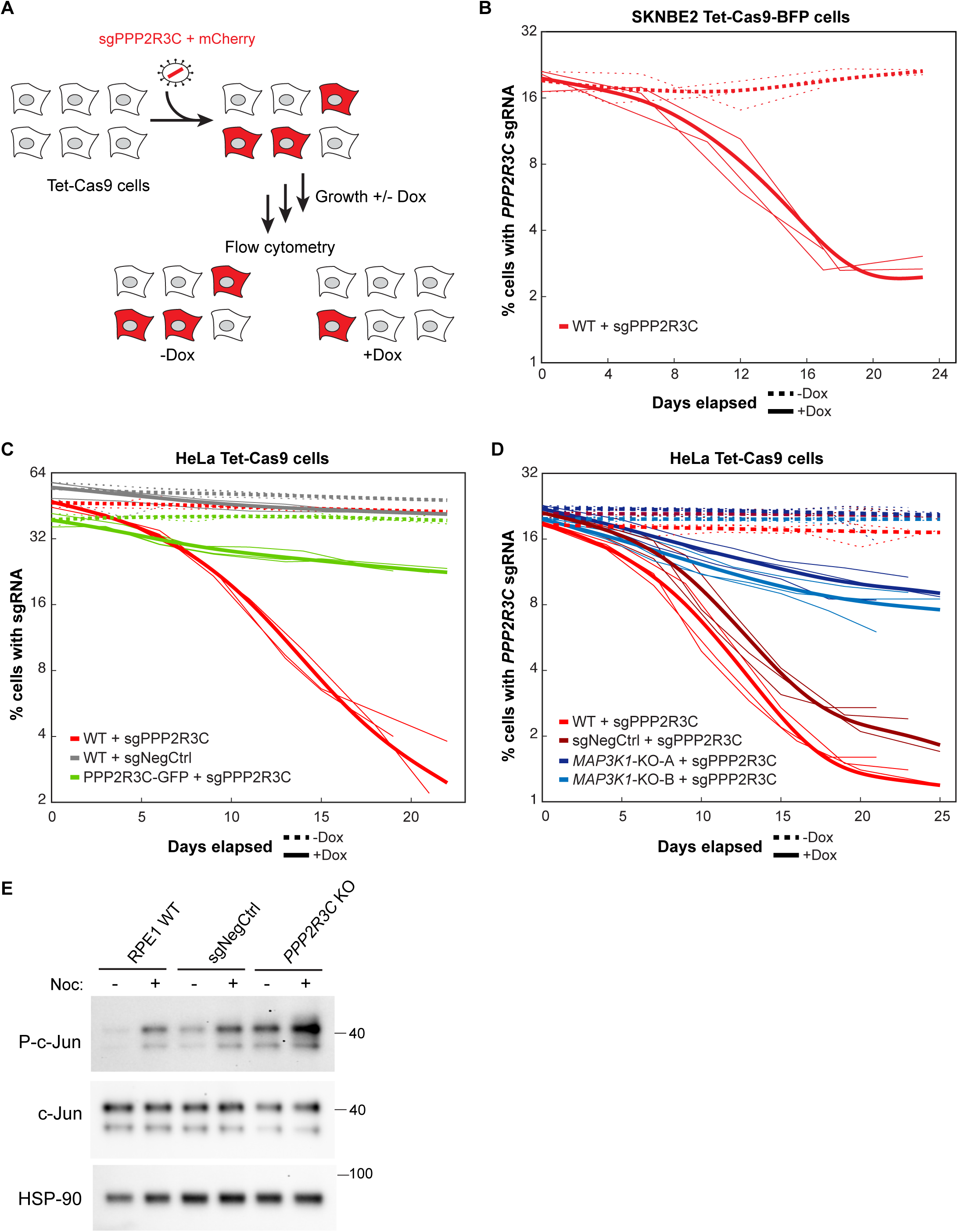
PPP2R3C promotes cell fitness by opposing kinase MAP3K1. **A)** Illustration of flow cytometry-based competitive growth assay in which cells expressing Cas9 are transduced with a lentiviral construct co-expressing an sgRNA targeting a gene of interest (such as *PPP2R3C*) and mCherry to mark transduced cells. Flow cytometry analysis at successive timepoints allows monitoring of the relative growth of mutant cells relative to untransduced wildtype cells. **B)** The percentage of mCherry-positive cells expressing sgRNAs targeting *PPP2R3C* is plotted over time for SKNBE2 Tet-Cas9 cells growth with or without doxycycline (Dox)-mediated Cas9 induction. Thin lines show *N* = 3 replicate experiments, and thick lines show smoothing spline curves fit to replicates. **C)** The percentage of mCherry-positive cells is plotted as in (**B**) for sgRNAs targeting *PPP2R3C* or a non-targeting control (sgNegCtrl) in wildtype HeLa Tet-Cas9 cells or HeLa Tet-Cas9 cells stably expressing a sgRNA-resistant *PPP2R3C-GFP* transgene. **D)** The percentage of mCherry-positive cells is plotted as in (**B**) for sgRNAs targeting PPP2R3C introduced into wildtype HeLa Tet-Cas9 cells, HeLa Tet-Cas9 cells expressing a non-targeting sgRNA, or clonal *MAP3K1* KO HeLa Tet-Cas9 cell lines. See also Figure S3A-B. **E)** Immunoblot analysis of phosphorylated c-Jun (P-c-Jun) and total c-Jun levels in whole cell lysates prepared from the indicated cells grown for 1 hr in the presence or absence of 2 μg/ml Nocodazole (Noc). HSP90 was used as a loading control. One of *N* = 3 replicate experiments.

While our analyses above revealed that *MAP3K1* expression levels are correlated with the severity of *PPP2R3C* KO phenotypes across cell lines, whether MAP3K1 has a causal role remained unclear. To address this key question, we asked if *PPP2R3C* KO growth defects would be rescued by knockout of *MAP3K1*. Here, we sought to use a more experimentally amenable cell line in which a *PPP2R3C* KO growth phenotype could be readily induced and measured. To this end, we made use of HeLa cells with inducible Cas9 expression that were recently used for systematic exploration of essential gene function.^58^ In these cells, we first established that *PPP2R3C* KO causes a fitness defect using the same flow cytometry-based assay as for SKNBE2 cells (Fig. 3C). Confirming the specificity of this phenotype, no depletion of mCherry-positive cells was seen for a non-targeting control sgRNA, and the depletion of *sgPPP2R3C* cells was rescued by an sgRNA-resistant PPP2R3C-GFP transgene (also confirming the functionality of this construct used above) (Fig. 3C). We next used CRISPR to prepare two *MAP3K1* KO clones, which grow at normal rates (Fig. S3A-B). Notably, when the mCherry-marked *PPP2R3C*-targeting sgRNA cassette was introduced into *MAP3K1* KO HeLa cells, the *PPP2R3C* KO growth phenotype observed above was strongly suppressed (Fig. 3D). To confirm these results, we stained cells for PPP2R3C at the end of experiment. When wildtype cells were transduced with *PPP2R3C*-targeting sgRNAs, virtually no cells lacking centriolar PPP2R3C could be detected after 18-25 days (only 4 out of 1882 cell examined; thus, the flow cytometry analysis likely underestimates the growth defect because most surviving Cherry-positive cells apparently failed to knock out *PPP2R3C*). By contrast, PPP2R3C-negative cells could be readily observed in *MAP3K1* KO backgrounds, confirming that *MAP3K1* KO rescues the *PPP2R3C* KO growth phenotype (Fig. S3C). Lastly, we note that it was difficult to assess centrosome function following inducible knockout of *PPP2R3C* in a wildtype background due to heterogenous CRISPR kinetics and strong selective pressure on KO cells. In apparent *PPP2R3C* KO cells (lacking PPP2R3C centriolar staining), we noted a disorganized microtubule cytoskeleton with elevated peri-nuclear microtubule signal but did not observe the severe centriole disintegration phenotype seen upon MAP3K1 overexpression (Fig. S3D). Taken together, these analyses indicate that growth defects caused by *PPP2R3C* KO are due to excess MAP3K1 kinase activity.

To test if MAP3K1 and PPP2R3C having opposing effects on kinase signaling, we assessed Jun phosphorylation, an established output of MAP3K1 activity.^57^ In wildtype RPE1 cells, we observed increased levels of phosphorylated Jun (P-Jun) upon microtubule depolymerization by nocodazole, as previously reported (Fig. 3E).^57^ Consistent with PPP2R3C-mediated phosphatase activity opposing MAP3K1, we found that basal and nocodazole-induced P-Jun levels were strongly increased in *PPP2R3C* KO cells (Fig. 3E). A similar increase in P-Jun levels was also seen in Tet-GFP-Map3k1 cells (Fig. S3E). Thus, the effects of PPP2R3C and MAP3K1 on cell proliferation are reflected in their opposing effects on JNK signaling.

### A pathogenic PPP2R3C variant is deficient in centriolar function

Following our identification of *Ppp2r3c* in a screen for ciliary signaling mutants, pathogenic variants in *PPP2R3C* were reported in patients with a developmental syndrome characterized by 46-XY complete gonadal dysgenesis, facial dysmorphism, and retinal degeneration.^9–11^ This myoectodermal gonadal dysgenesis (MEGD) syndrome (OMIM #618419) is a recessive disorder caused by mutations at conserved PPP2R3C residues L193S, F350S, or L103P (Fig. 4A).^10,11^ However, the mechanistic basis by which PPP2R3C mutations cause developmental defects remains unknown. Strikingly, a similar form of gonadal dysgenesis is also reported in individuals with heterozygous variants in *MAP3K1* that are proposed to be dominant gain-of-function mutations (OMIM #613762).^12–15^ Thus, loss-of-function variants in *PPP2R3C* and gain-of-function variants in *MAP3K1* may have a shared impact on gonadal development, further supporting the opposing functional roles we identified above for these proteins.

**Figure 4.**
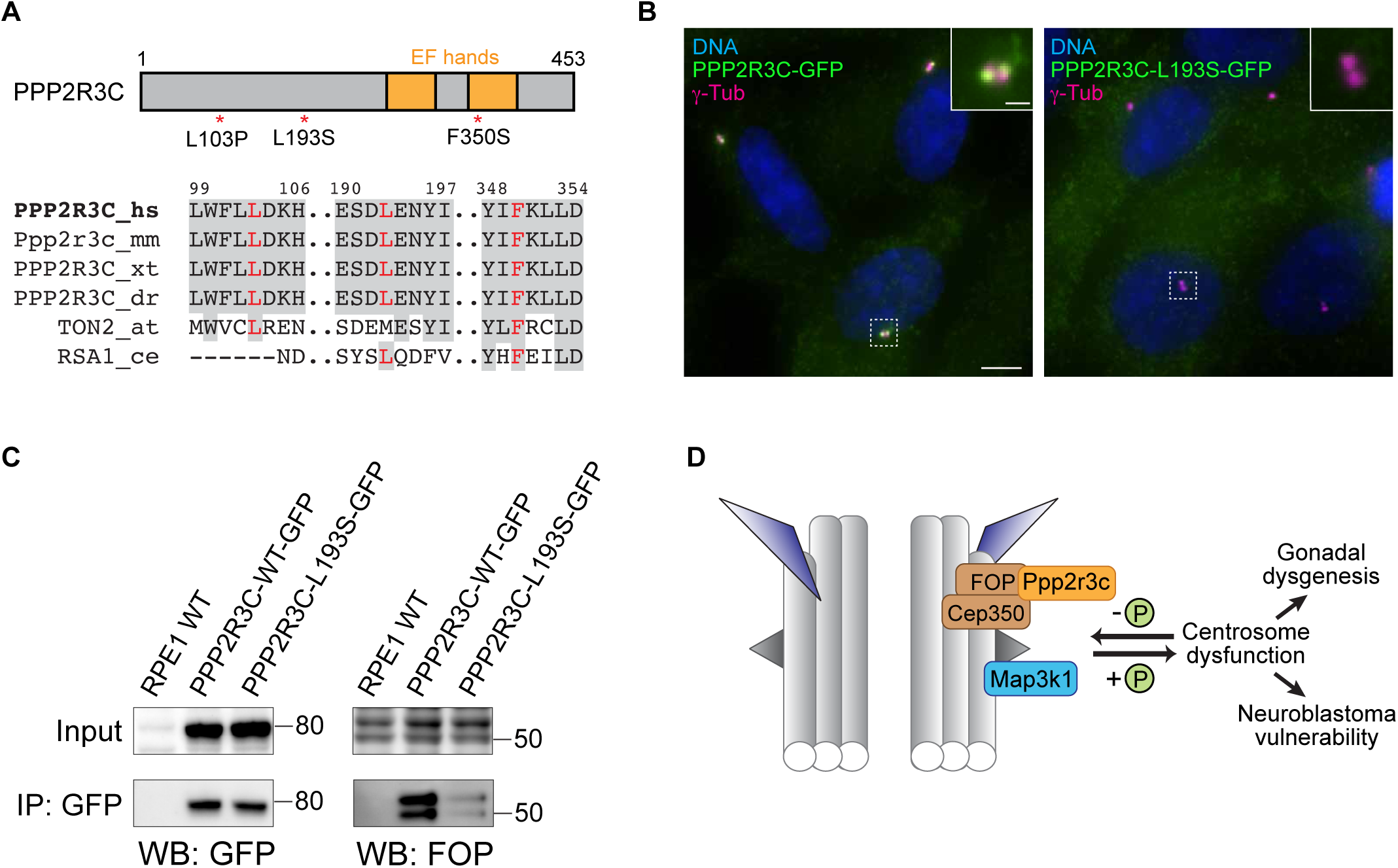
A human pathogenic variant in PPP2R3C impairs its centriolar function. **A)** Top: Illustration of PPP2R3C highlighting two EF hand domains and localization of three pathogenic variants observed in individuals with myo-ectodermal gonadal dysgenesis (MEGD) syndrome. Bottom: alignment of PPP2R3C sequence showing human (hs), mouse (mm), frog (*Xenopus tropicalis*; xt), zebrafish (*Danio rerio*; dr), and plant (*Arabidopsis thaliana*; at) homologs. Residues mutated in MEGD are shown in red, and conserved residues are highlighted gray. **B)** Immunofluorescence microscopy analysis of wildtype and L193S forms PPP2R3C-GFP stably expressed in RPE1 cells. Centrosomes are marked by ψ-tubulin; ψ-tub). One of *N* = 3 representative experiments. Scale bar: 5 μm; inset: 1 μm. **C)** Proteins captured via anti-GFP immunoprecipitation from wildtype RPE1 cells and from RPE1 cells stably expressing wildtype or L193S forms of PPP2R3C-GFP were detected by immunoblotting, as indicated. One of *N* = 3 representative experiments. **D)** Model for PPP2R3C and MAP3K1 function, indicating association of PPP2R3C with FOP and CEP350 and its localization at the centriole distal torus. MAP3K1 kinase and PP2A-PPP2R3C phosphatase have opposing roles, with excess MAP3K1 or loss of PPP2R3C promoting gonadal dysgenesis and fitness defects, including in cancers such as neuroblastoma that are characterized by high *MAP3K1* expression.

To test if syndromic mutations in PPP2R3C affect the centriolar activity of this phosphatase, we stably expressed a PPP2R3C-L193S-GFP variant in wildtype RPE1 cells, as this mutation has been observed in the largest number of MEGD individuals to date.^11^ Notably, while wildtype PPP2R3C-GFP was readily detected at mother and daughter centrioles, we did not observe centriolar localization for the L193S mutant even though it was expressed in RPE1 cells at similar levels as a wildtype transgene (Fig. 4B-C). Similarly impaired localization was also observed for this variant in HeLa cells (Fig. S4A). To investigate the basis for this altered localization, we assessed the impact of the L193S mutation on binding to centriolar protein FOP. Notably, when wildtype and L193S forms of PPP2R3C-GFP were immunoprecipitated from RPE1 cells, PPP2R3C-L193S exhibited greatly diminished binding to FOP compared to wildtype PPP2R3C (Fig. 4C). Thus, our findings collectively indicate that MEGD-associated PPP2R3C variants may impair the centriolar function of PPP2R3C phosphatase through an impaired localization, leading to a similar pathology as gain-of-function mutations in MAP3K1 kinase (Fig. 4D).

## Discussion

Here we identify the PP2A regulatory subunit PPP2R3C as a centriolar protein that acts with FOP and CEP350 to counteract signaling by MAP3K1. Imbalance of this phospho-regulatory module strongly inhibits cell growth, with excess MAP3K1 activity disrupting centrosomal microtubule nucleation, cilium formation, and centriole structural integrity. The importance of balanced activities of PPP2R3C and MAP3K1 is further illustrated by pathogenic variants in these proteins that cause gonadal dysgenesis, and we demonstrate that a disease-associated PPP2R3C mutation impairs its centriolar localization and binding to FOP. Thus, our study provides new insights into centriole function, illuminates pathologies caused by defective centrosomal proteins, and reveals the power of functional genomics to provide insight into uncharacterized gene functions.

While we initially identified PPP2R3C in a screen for Hh signaling regulators, the central model we proposed arose from our analysis of the DepMap dataset. A powerful feature of this resource is the ability to generate unbiased functional predictions for poorly characterized genes. We show that these predictions can emerge not only from co-essentiality analyses^46,48–51^, but also from integrating essentiality data with gene expression. Having validated the accuracy of these predictions, we believe such bioinformatic approaches will find broad utility as the scope of available datasets continues to grow.

A key conclusion from our study is that PPP2R3C is a centriolar protein that opposes MAP3K1 activity. Beyond the findings presented here, a centrosomal role for PP2A-PPP2R3C is further supported by the centrosomal localization of catalytic subunit PPP2CB reported in a recent large-scale study^37^ and by a systematic analysis of essential genes.^58^ In the latter report, Funk et al. used multi-parametric microscopy to characterize and cluster essential genes, with *PPP2R3C* falling in a shared cluster with centrosomal genes *PLK4*, *SASS6*, *CEP350*, *PCM1*, and *STIL*.^58^ For MAP3K1, we find that over-expression of this kinase strongly impairs centrosome function and produces a striking rapid disintegration of centrioles. Thus, PPP2R3C inactivation and MAP3K1 over-expression phenotypes converge on centrosome dysfunction, reflecting the centriolar localizations we observed for these proteins.

Our study also provides new insight into key centriolar proteins FOP and CEP350 by linking these proteins to PPP2R3C. Based on the co-essentiality, co-localization, and co-immunoprecipitation of these proteins, we propose that one essential function of FOP and CEP350 is to support PPP2R3C in counteracting MAP3K1. In prior studies, FOP and CEP350 have been found to regulate ciliogenesis, microtubule nucleation, centriole length, and centriole stability.^52,54,59,60^ Thus, FOP and CEP350 likely regulate diverse centrosomal functions through PPP2R3C as well as by PPP2R3C-independent mechanisms.

Our finding of a novel PPP2R3C-MAP3K1 phospho-regulatory module at centrosomes lays the foundation for future studies to elucidate the substrates targeted by these proteins. While we find that Jun phosphorylation is jointly regulated by PPP2R3C and MAP3K1, other substrates of JNK signaling and/or outputs of MAP3K1 may contribute to centrosome dysfunction induced by MAP3K1. Given the rapid loss of centriole integrity triggered by MAP3K1 overexpression, defining these substrates may provide valuable insights into how centriole structure is maintained. Additionally, it will be important to define conditions under which the balance of these enzymes is physiologically modulated to regulate centrosome function. Microtubule stress may be one such condition, as P-Jun levels are strongly elevated upon nocodazole treatment. An additional possibility is that MAP3K1 and PPP2R3C may contribute to centrosome inactivation and centriole elimination, such as during muscle differentiation or oocyte production.^61^ With the tools and phenotypes defined here, these will be exciting areas for future study.

The physiologic importance of PPP2R3C-MAP3K1-mediated phospho-regulation is illustrated by the finding that both proteins are mutated in gonadal dysgenesis syndromes. While *PPP2R3C* and *MAP3K1* have been recognized among several gonadal dysgenesis genes, we establish here a previously unknown functional connection between them. Interestingly, their opposing functional roles are reflected in the fact that contrasting types of mutations and modes of inheritance that cause disease (recessive, inactivating PPP2R3C variants and dominant, activating MAP3K1 variants). A key question for future study will be to determine why select tissues and processes are most strongly impacted by these mutations. Given that PPP2R3C-L193S fails to bind FOP or localize to centrioles, a compelling model for further investigation is that centrosome dysfunction may underlie these disorders.

Beyond their relevance to developmental syndromes, PPP2R3C and MAP3K1 may also provide insight into cancer biology. Neuroblastomas and some blood cancers (e.g. B-cell leukemias) commonly exhibit high *MAP3K1* expression and a strong dependence on *PPP2R3C*.^47^ Consistent with this trend, in the DT40 avian B cell leukemia cell line, *FOP* is essential for viability, and microtubule stress induces apoptosis in a MAP3K1-dependent manner.^62,63^ Thus, PPP2R3C may represent a selective vulnerability in certain cancers that could be therapeutically exploited, potentially via microtubule-targeting drugs that activate MAP3K1.

While our studies have focused on mammalian cells, *PPP2R3C* homologs are present in diverse species. In *C. elegans*, RSA-1 is homologous to PPP2R3C (as well as PPP2R3A and PPP2R3B) and has key roles in microtubule nucleation and mitotic spindle formation.^35^ However, *C. elegans* lacks known homologs of *FOP* or *CEP350*, and RSA-1 functions instead through association with PCM protein SPD-5. In plants, *TON2* is a *PPP2R3C* homolog and a key regulator of microtubule organization in interphase and at mitotic entry.^64^ Notably, TON1 is also required for plant microtubule organization and has partial homology to FOP.^65^ TON1 further interacts with proteins bearing a TRM motif, a sequence element that is also found in human CEP350.^66^ Connecting these lines of evidence, a recent study found that a PP2A-TON2-TON1-TRM protein complex is needed for microtubule organization in plants.^67^ We therefore propose that a human PP2A-PPP2R3C-FOP-CEP350 complex is a functional ortholog of this complex that has become specialized for centrosomal microtubule nucleation and incorporated into centrioles. Physical association among these human proteins is supported by interactome studies and by the co-precipitation we observe between PPP2R3C and FOP.^37,68,69^ Thus, PPP2R3C is part of a phospho-regulatory module that is conserved across divergent modes of microtubule organization and that is required in human cells is to restrain MAP3K1 signaling, thereby regulating centrosome function in development and disease.

## Acknowledgements

The authors acknowledge Iain Cheeseman for sharing HeLa Tet-Cas9 cells, Ben Turk for helpful discussions, the Yale Science Hill FACS facility, and funding from the National Institutes of Health (R35GM137956 to D.K.B.), a National Science Foundation Graduate Research Fellowship (to C.S.B.), the Swiss State Secretariat for Education, Research and Innovation (SERI) (MB22.00075 to P.G.), and the Swiss National Science Foundation (SNSF) (310030_205087 to P.G. and V.H.).

## Author Contributions

A.K.G. and D.K.B. conceived the project. A.K.G., L.K.S., A.R.R., and C.S. conducted experiments and analyzed results. D.K.B. and V.H. and P.G. supervised the project. A.K.G. and D.K.B. wrote the manuscript with input from all authors.

## Supplementary Information

Supplementary Figures 1-4

## Methods

### Resource Availability

#### Lead Contact

Further information and requests for resources and reagents should be directed to and will be fulfilled by the lead contact, David Breslow.

#### Materials Availability

All unique reagents generated in thus study are available from the lead contact upon request.

#### Data and code availability

Raw data not included in this article are available from the corresponding author on request. Code for U-ExM analysis is available at https://github.com/CentrioleLab/pickCentrioleDim.

### Experimental model and subject details

#### Cell Culture

RPE1-hTERT (RPE1), SKNBE2, and HEK 293T cells were obtained from ATCC. HeLa cells with inducible Cas9 were a gift from Iain Cheeseman (Whitehead Institute of MIT). All cell lines were grown in a 37°C humidified chamber with 5% CO2. Cell lines were routinely checked for mycoplasma contamination using MycoAlert Plus detection kit (Lonza). HEK293T cells were grown in DMEM high-glucose medium (Gibco) supplemented with 10% fetal bovine serum (FBS; HyClone), 100 units/mL Penicillin (Gibco), 100 mg/mL Streptomycin (Gibco), 2 mM Glutamine (Gibco) and 1 mM sodium pyruvate (Gibco). RPE1-hTERT cells were grown in DMEM/F12 medium (Gibco) supplemented with 10% FBS (Gemini Bio), 100 units/mL Penicillin (Gibco), 100 mg/mL Streptomycin (Gibco), 2 mM Glutamine (Gibco). HeLa cells were grown in HEK293T medium except 10% tetracycline-free FBS (Takara Bio) was used. SKNBE2 cells were grown in Eagle’s Minimum Essential Medium (EMEM; ATCC) with the same supplement as used for HeLa cell medium. To induce ciliogenesis, RPE1-hTERT cells were serum-starved in medium containing 0.2% FBS for 48 hrs. For inducible gene expression, cells were treated with 1 μg/ml doxycycline (Fisher Scientific). To induce microtubule depolymerization, cells were treated with 2 μg/ml Nocodazole for 1 hr at 37°C.

### Method details

#### DNA cloning

CRISPR sgRNAs were cloned either into pMCB320 (Addgene #89359, gift from Michael Bassik) or pKHH030 (Addgene #89358, gift from Michael Bassik). For pMCB320, sgRNAs were cloned from annealed oligonucleotides inserted between BstxI and BlpI restriction sites. For pKHH030, sgRNAs were similarly inserted between BbsI and Bpu1102I sites. For increased CRISPR efficiency, a pMCB320-sgPPP2R3C.1-sgPPP2R3C.2 double-sgRNA construct was used by cloning sgPPP2R3C.A into pKHH030 and the sub-cloning the XhoI and BamHI-flanked sgPPP2R3C.A cassette from pKHH030 into pMCB320_sgPPP2R3C.B digested with the same enzymes. All sgRNA sequences are listed in the Key Resource Table. DNA fragments containing MAP3K1 (MEKK1) or kinase-dead MAP3K1 were excised from pCDNA-MEKK1 plasmids (Addgene #12181 and #12180, respectively; gift from Gary Johnson) by DraI and EcoRV digestion. Resulting fragments were cloned into pENTR1A using the same restriction sites. Subsequently, Gateway cloning was used to transfer MAP3K1 cDNA sequences into PB-Tet-LAP-DEST (a plasmid produced by Gibson assembly from a piggyBac Tet-Cas9 plasmid kindly provided by Iain Cheeseman [MIT]), yielding PB-Tet-LAP-MAP3K1 plasmids suitable for PiggyBac transposition (LAP = GFP-TEV-S-tag). Human *PPP2R3C* cDNA was obtained from Horizon Discovery and cloned into pDONR221 by Gateway cloning. Silent mutations for sgRNA resistance and/or to encode the L193S mutation were introduced by gene synthesis and Gibson assembly. Stable expression of PPP2R3C-GFP was achieved by replacement of Rab34 with PPP2R3C-GFP in pCW-Pgk-Rab34_hPgk-miRFP-670-Centrin2^70^ (used in HeLa cells) and by replacement of Rab8 with PPP2R3C-GFP in pCW-Pgk-mScarlet-Rab8_hPgk-NeoR^71^ (used in RPE1 cells). For tetracycline-inducible expression of Cas9-BFP, pCW-Tet-Cas9-BFP was prepared by Gibson assembly from pCW-Cas9 (Addgene #50661).

#### Lentivirus production

Lentiviral particles were produced by co-transfection of HEK293T cells with a lentiviral vector and packaging plasmids (pCMV-VSVG, pRSV-Rev, and pMDLg/RRE for sgRNA plasmids and pCMV-VSVG and pCMV-ΔR-8.91 for all other constructs). After transfection with polyethyleneimine (linear, MW ∼25,000), virus-containing supernatant was collected, filtered through a 0.45 µm polyethersulfone filter and used directly or concentrated 3-fold with Lenti-X Concentrator (Clontech).

#### Production of stable cell lines

RPE1, HeLa, and SKNBE2 cell lines were modified for expression of transgenes or sgRNAs by lentiviral transduction or piggyBac-mediated transposition. SKNBE2 cells with inducible Cas9 were prepared by transduction with pCW-Tet-Cas9-BFP. For CRISPR KO in RPE1 cells, sgRNAs were lentivirally transduced into RPE1 cells stable expressing Cas9-Blast.^71^ Following puromycin selection for sgRNA-expressing cells, clonal cell lines were isolated by FACS sorting on a FACS Aria III instrument (BD Scientific). HeLa *MAP3K1* KO cells were prepared by lentiviral transduction with pKHH030-sgMAP3K1-A, selected with puromycin, treated with doxycycline to induce Cas9, and FACS sorted to isolate clonal cell lines. All KO cell lines were validated by amplification and sequencing of genomic DNA (see Key Resource Table for primer sequences), as well as by immunofluorescent staining for the target gene product, where feasible. HeLa cells expressing sgRNA-resistant PPP2R3C-GFP were obtained by lentivirally transducing HeLa Tet-Cas9 cells with pCW-Pgk-PPP2R3C-GFP_hPgk-miRFP-670-Centrin2 and FACS sorting of a polyclonal pool of miRFP-670-positive cells. RPE1 cells expressing PPP2R3C-GFP and PPP2R3C-L193S-GFP were obtained by lentivirally transducing RPE Cas9-Blast cells with pCW-Pgk-PPP2R3C-(L193S)-GFP_hPgk-NeoR, followed by G418 selection to obtain polyclonal pools of PPP2R3C-GFP-expressing cells.

For piggyBac transposition, RPE1 cells were transfected with PB-Tet-LAP-MAP3K1 and PB-Transposase plasmid (kindly provided by N. Dimitrova) using X-tremeGENE9 reagent (Roche) and after 48 hours cells were selected with G418. Selected cells were then single-cell sorted using a FACS Aria III (BD Scientific) instrument to isolate clonal cell lines that were validated by assessing doxycycline-induced MAP3K1 expression.

#### Immunofluorescence staining and microscopy

Cells were seeded on acid washed #1.5 coverslips in 24-well plates and fixed in either 4% paraformaldehyde for 10 minutes, -20°C methanol, or both in succession. For microtubule regrowth assays, coverslips were placed on ice for 30 min, followed by transfer to room temperature for the indicated times to allow microtubule formation. Coverslips were washed three times with PBS and permeabilized with PBS containing 0.1% Triton X-100 for ten minutes. Coverslips were washed twice with PBS, blocked in PBS containing 3% BSA and 5% Normal Donkey Serum for twenty minutes. Blocked coverslips were incubated with primary antibodies for 40 minutes. After incubation with primary antibodies (see Key Resource Table), coverslips were washed five times with PBS containing 3% BSA and incubated with secondary antibodies. After 30 minutes of incubation, coverslips were washed and incubated with Hoechst 33258 DNA dye for 5 minutes, washed, and mounted on glass slides using Fluoromount G (Electron Microscopy Sciences). Stained coverslips were imaged using a Nikon Eclipse Ti-2 widefield microscope equipped with a 60X PlanApo oil objective (NA 1.40; Nikon Instruments), Prime BSI camera (Photometrics), and LED light source (Lumencor SOLA-V-NIR). Samples were imaged at room temperature using Nikon Elements software and analyzed in Fiji/ImageJ.

#### Ultrastructure expansion microscopy U-ExM

U-ExM was performed as described previously.^53^ RPE1 cells seeded on 12 mm coverslips were fixed in a 2% acrylamide / 1.4% formaldehyde solution in PBS for 3 hrs at 37°C, except for the endogenous staining of PPP2R3C protein, where the cells were first fixed with methanol for 10 minutes at -20°C. Fixed cells were next embedded in a gel matrix formed by combining Monomer Solution (19% sodium acrylate, 0.1% bis-acrylamide, and 10% acrylamide) with TEMED and APS (final concentration of 0.5%) for 1 hr at 37°C in a humidified chamber. Samples were incubated for 1.5 hr at 95°C in Denaturation Buffer (200 mM SDS, 200 mM NaCl, 50 mM Tris, pH 9). Finally, gels were placed in distilled water for 1 hr, and the expansion factor of the gel was measured. Gels were then punched in small circles and stored in 50% glycerol until immunostaining.

For immunostaining, gels were washed three times with PBS and then incubated overnight at 4°C with primary antibodies diluted in PBS supplemented with 2% BSA. After 3 washes in PBS-Tween 0.2%, gels were incubated in secondary antibodies for 2 hrs at 37°C. Finally, gels were washed in PBS-Tween 0.2% 3 times and then placed in distilled water for final expansion.

To image U-ExM samples, gels were placed on poly-D-Lysine-coated 24 mm coverslips to avoid sample drift. Image acquisition was done using a Leica Thunder DMi8 microscope or a Leica Stellaris 8 Falcon. In both cases, a 63×/1.4 NA oil objective was used. Images were deconvolved using Leica Thunder SVCC (Thunder DMi8 images) software or the Lightning mode at highest resolution (Stellaris 8 Falcon images). ‘Water’ was chosen as the mounting medium in both deconvolution softwares. Images were processed with ImageJ (Fiji).

#### Growth competitions and flow cytometry

For growth competition experiments, the indicated HeLa or SKNBE2 cell lines were transduced at moderate multiplicity of infection with a double-sgRNA construct targeting *PPP2R3C* (or a non-targeting control) and co-expressing mCherry. Following transduction, cells were split into doxycycline-treated or untreated plates and cultured for >18-25 days, with flow cytometry analysis conducted at each timepoint using a Fortessa X-20 or FACS Aria III instrument (Becton Dickinson). The fraction of mCherry-positive cells was then analyzed over time for N = 3 replicate experiments. For SKNBE2 cells, quantification of doxycycline-induced cells was restricted to Cas9-BFP-positive cells, as a modest fraction of cells were refractory to Cas9-BFP induction. Figures show a curve fitted to the combined data from replicate experiments using the smoothing spline method from the curve fitting toolbox of Matlab software (The Mathworks, Inc.).

For cell cycle analysis by propidium iodide staining, cells were seeded at sub-confluent density and grown for 24 hr prior to collection. Cells were harvested via trypsinization, washed in PBS, resuspended in 1 ml PBS, and then added dropwise to 4 ml ice cold ethanol while vortexing. Cells were fixed at -20°C for >30 min, collected by centrifugation and resuspended in PBS containing 0.05% Triton X -100, 10 μg/ml RNAse, and 20 μg/ml propidium iodide . After filtering through a 35 μm strainer cap, cells were incubated for 30 min at room temperature and analyzed on BD FACS Aria III or Fortessa X-20 instruments (Becton Dickinson). The resulting data were analyzed using FlowJo software (Becton Dickinson).

#### Immunoprecipitation and immunoblotting

For immunoprecipitations, cells were seeded in 10 cm dishes, cultured for 24 hr, and then collected by trypsinization, washed twice in PBS, pelleted, and flash frozen in liquid nitrogen. Frozen cells were then lysed in CoIP buffer (50 mM Tris pH 7.4, 150 mM NaCl, 1% Tx-100, 0.5 mM DTT, and protease inhibitors) on ice for 15 min. Lysate was then centrifuged at 20,000 x *g* at 4°C, and the supernatant was collected into a fresh tube. Protein concentrations were quantified using Bio-Rad Protein Assay and diluted to equal concentrations. GFP-Trap beads (ChromoTek) pre-washed with CoIP buffer were then incubated for 75 minutes with 100 μl cell lysate at 4°C. After incubation, beads were collected by centrifugation at 1000 rpm for 30 seconds, washed six times with Co-IP buffer with protease inhibitors, and bound proteins were eluted by boiling in 1X NuPage LDS buffer.

To analyze protein levels in whole cell lysate, cells were plated in 6-well plates, cultured for 24 hr, and then lysed by scraping on ice in ice-cold RIPA buffer (50 mM Tris pH 7.4, 150 mM NaCl, 2% Igepal CA-630, 0.25% sodium deoxycholate, 0.5 mM DTT, protease inhibitors). After incubation on ice for 10 min, lysates were centrifuged at 20,000 x *g* for 10 min at 4°C and supernatants collected. Protein concentrations were determined using Bio-Rad Protein Assay reagent, and lysates diluted as needed to normalize protein concentrations.

For SDS-PAGE and immunoblotting, whole cell lysates or immunoprecipitated samples were separated on NuPAGE 4-12% Bis-Tris protein gels (Invitrogen) in MOPS buffer and then transferred to PVDF membrane (Millipore Sigma). Membranes were blocked with 1:1 PBS/SeaBlock (Thermo Fisher) for 40 min at room temperature. Blocked membranes were washed with TBST for 5 minutes and incubated with primary antibodies either for 1.5 hr at room temperature (anti-HSP90, anti-Jun, and anti-P-c-Jun) or 16 hr at 4°C (anti-Fop and anti-GFP). Membranes were then washed with TBST three times, incubated with Protein A-HRP for 30 min at room temperature, washed again three times in TBST, and developed using Clarity Western ECL Substrate (Bio-Rad). Membranes were imaged on a ChemiDoc Touch imager (Bio-Rad) according to manufacturer’s instructions.

### Quantification and statistical analysis

Analysis of cancer cell line growth data in the DepMap (Dependency Map) data set was carried out using the CRISPR gene effect data and CCLE gene expression data in the 2022Q2 release.^47^ Pearson correlation coefficients were calculated in Matlab (Mathworks, Inc.) between the gene effect profile of *PPP2R3C* across all cell lines and the corresponding gene effect profiles for all other genes. To assess the relationship between gene expression levels and PPP2R3C gene effect, correlations were calculated using the cell lines present in both the gene effect and gene expression datasets. Specifically, correlation coefficients were calculated as above between the gene effect profile of *PPP2R3C* and each gene’s profile of gene expression across cell lines. To examine cancer-type-specific effects, data were grouped and averaged by cancer lineage subtype reported in the DepMap dataset (note: only cancer types represented by at least five cell lines were included for analysis). Comparison of *PPP2R3C* gene effect data across cell lines to *IFT88* gene effect data is based on data in the 2022Q4 release.^72^

To quantify the localization of proteins analyzed by U-ExM, the plugin pickCentrioleDim was used (available at https://github.com/CentrioleLab/pickCentrioleDim). This plugin measures the distance between the 50% peak of intensity of the protein of interest relative to tubulin, indicating both the length and the width of the staining patterns. The final reported distances and dimensions were modified according to the expansion factor of each experiment to indicate the non-expanded staining patterns.

All statistical tests were based on two-tailed, unequal variance t-test, with the exception of centriole number analysis in Fig. 2H. For comparison of centriole numbers, two-way ANOVA with Dunnett multiple comparison correction (single pool variance) was performed.

## KEY RESOURCES TABLE

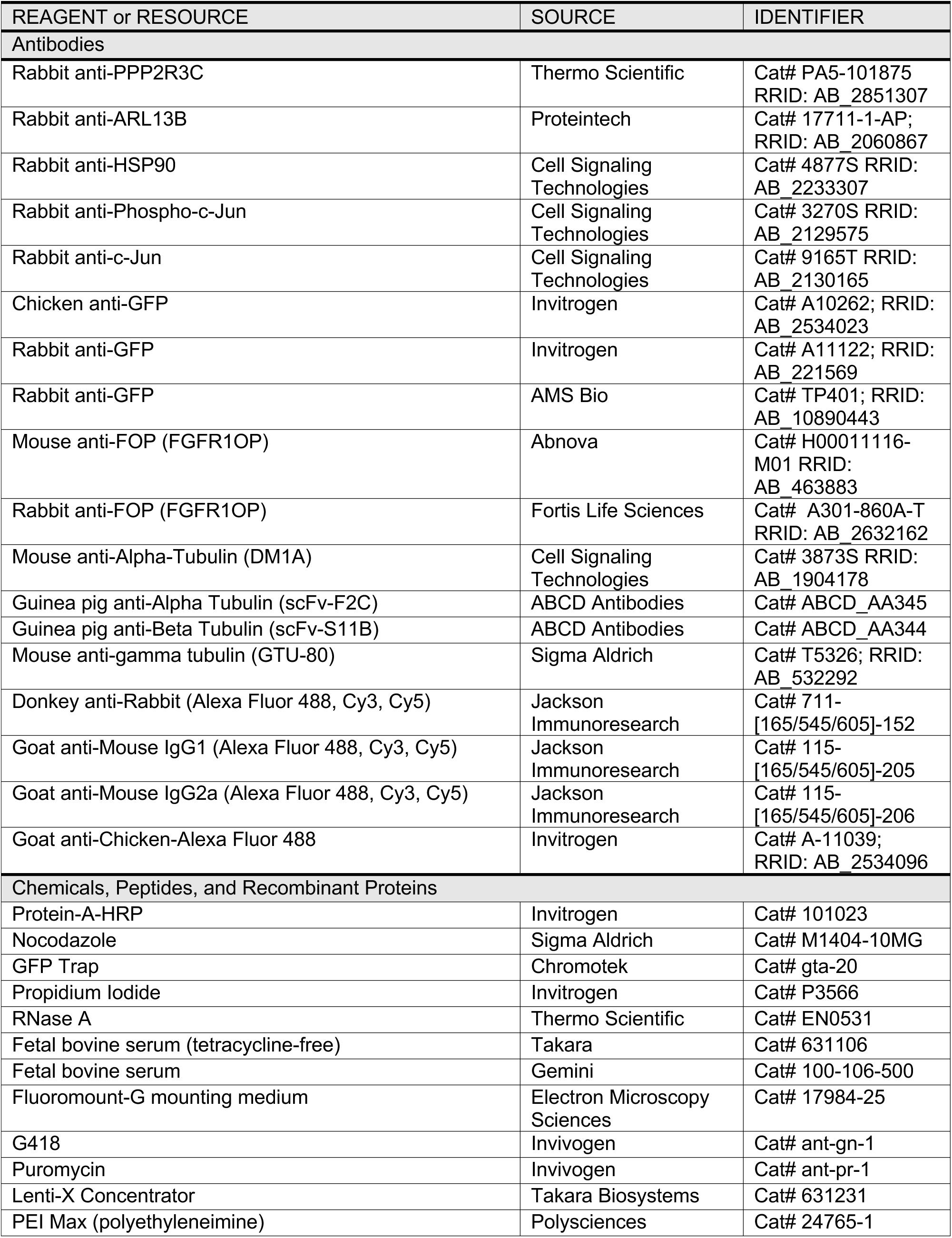
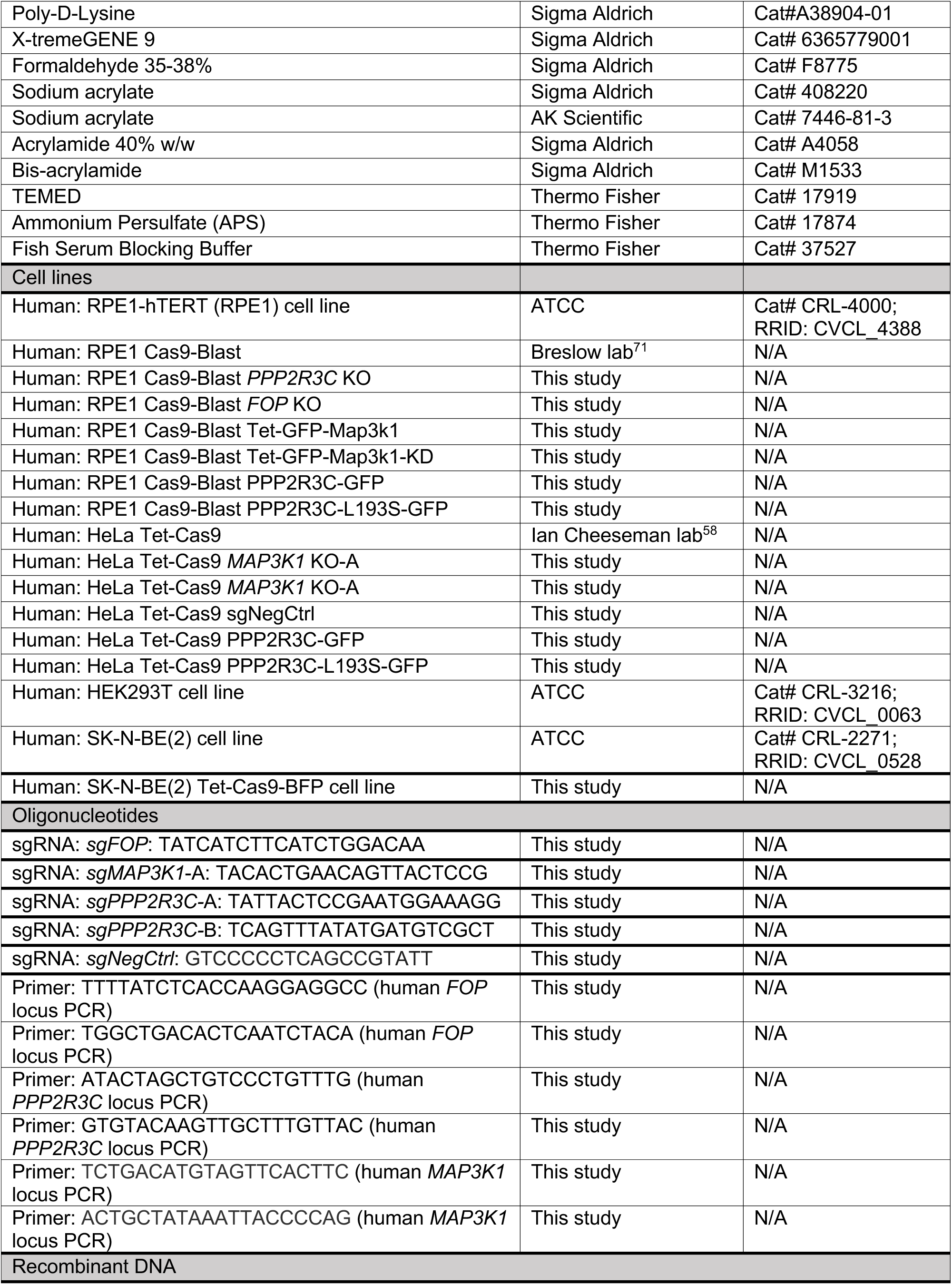
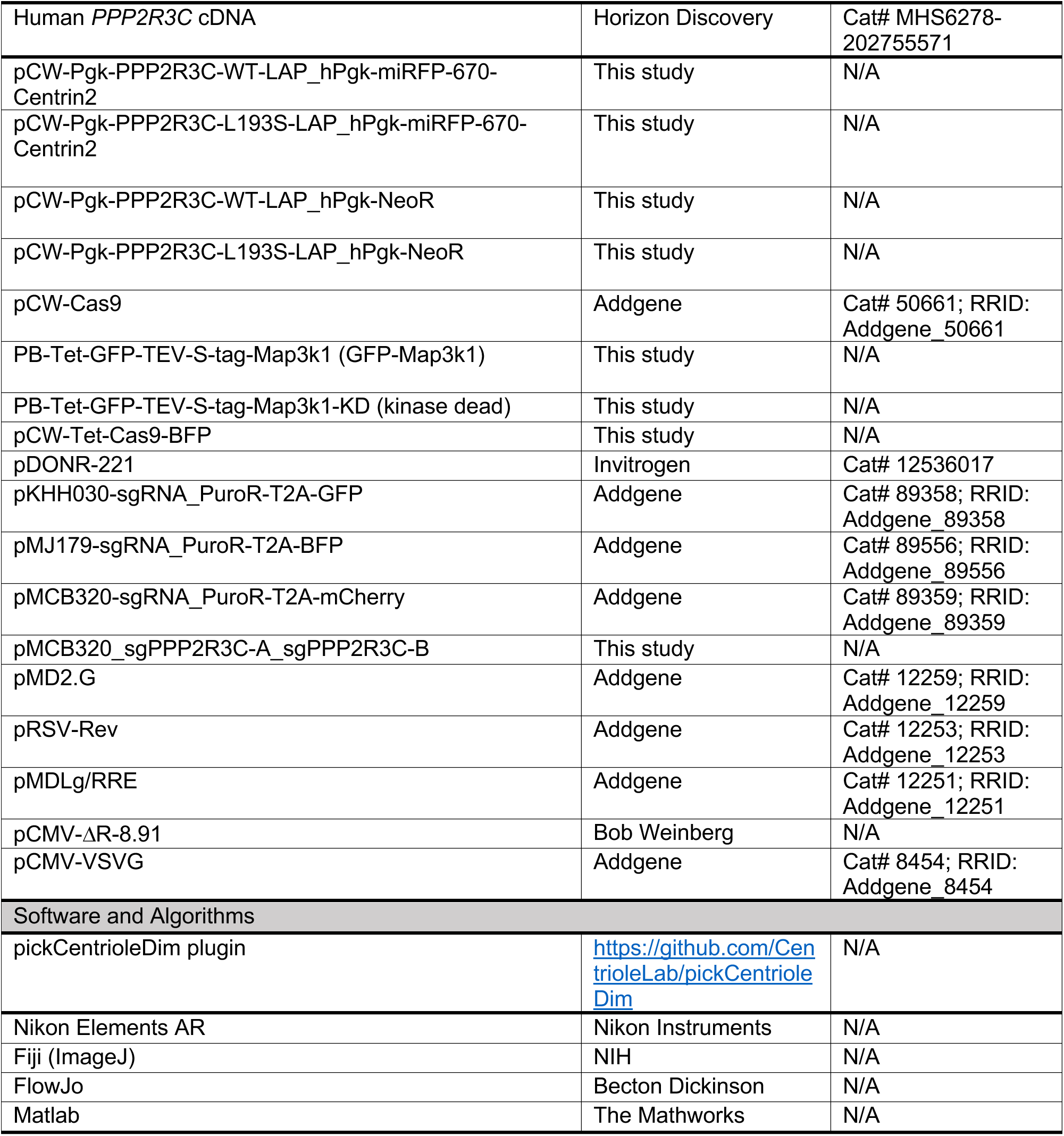

