## Supplementary Materials for "A disease-associated PPP2R3C-MAP3K1 phospho-regulatory module controls centrosome function"

**Supplementary Materials for Ganga et al.**

Supplementary Figures 1-4

**Figure S1**

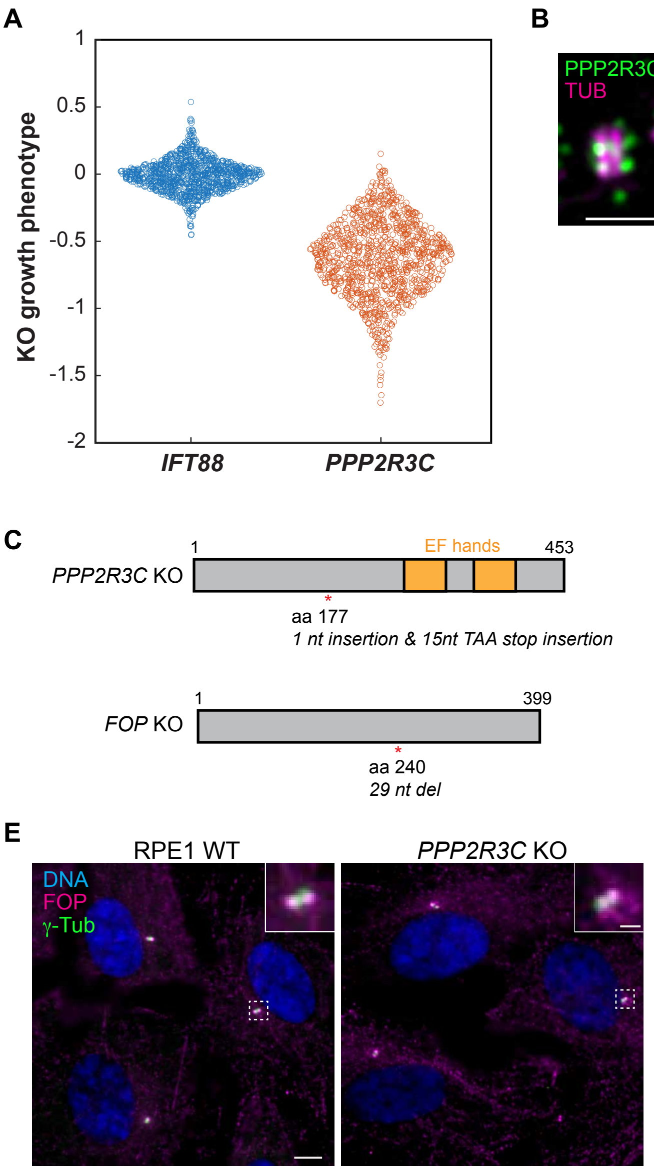
A
KO growth phenotype
1
0.5
0
-0.5
-1
-1.5
-2
IFT88
PPP2R3C
B
PPP2R3C
TUB
TUB
PPP2R3C
C
PPP2R3C KO
1
EF hands
453
aa 177
1 nt insertion & 15nt TAA stop insertion
FOP KO
1
399
aa 240
29 nt del
D
RPE1 WT
DNA
FOP
γ-Tub
FOP KO
E
RPE1 WT
PPP2R3C KO
DNA
FOP
γ-Tub

**Figure S1. Functional characterization of PPP2R3C**

**A)** Growth phenotypes for CRISPR targeting of *IFT88* and *PPP2R3C* are shown for cancer cells included in the DepMap dataset. **B)** Representative U-ExM image of wildtype RPE1 cells stained for endogenous PPP2R3C and tubulin (TUB). Scale bar: 1 μm. **C)** Illustration of mutations validated by Sanger sequencing of RPE1 *PPP2R3C* and *FOP* KO cell lines generated by CRISPR. **D)** Immunofluorescence analysis of FOP and γ-tubulin in wildtype RPE1 cells and *FOP* KO cells. Scale bar: 5 μm; insets: 1 μm. **E)** Immunofluorescence analysis of FOP and centrosomes (marked by γ-tubulin; γ-tub) in wildtype RPE1 cells and *PPP2R3C* KO cells. Scale bar: 5 μm; insets: 1 μm.

### Figure S2

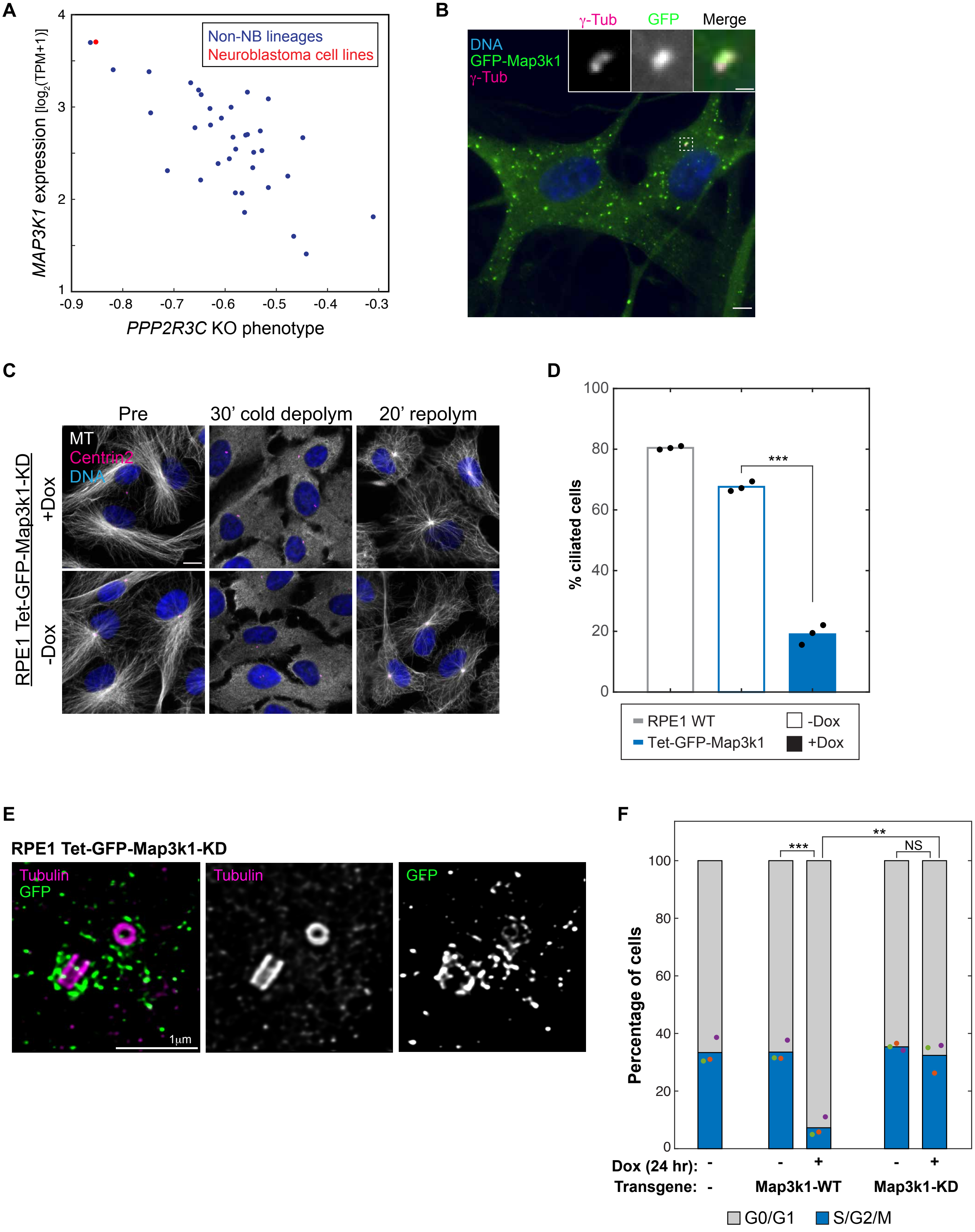
A
Non-NB lineages
Neuroblastoma cell lines
MAP3K1 expression [log2(TPM+1)]
PPP2R3C KO phenotype
B
γ-Tub
GFP
Merge
DNA
GFP-Map3k1
γ-Tub
C
Pre
30' cold depolym
20' repolym
MT
Centrin2
DNA
+Dox
-Dox
RPE1 Tet-GFP-Map3k1-KD
D
% ciliated cells
RPE1 WT
Tet-GFP-Map3k1
-Dox
+Dox
E
RPE1 Tet-GFP-Map3k1-KD
Tubulin
GFP
1μm
F
Percentage of cells
Dox (24 hr):
Transgene:
Map3k1-WT
Map3k1-KD
G0/G1
S/G2/M
NS

**Figure S2.** ***MAP3K1*** **is associated with** ***PPP2R3C*** **mutant phenotype and centrosome dysfunction**

**A)** Relationship between MAP3K1 expression and PPP2R3C KO phenotype in the DepMap dataset, showing mean values for each cancer lineage subtype represented N ≥ 5 cell lines. Neuroblastoma cells are highlighted in red. **B)** Immunofluorescence microscopy showing localization of GFP-Map3k1 relative to centrioles (marked by γ-tub) in RPE1 cells expressing doxycycline (Dox)-inducible GFP-Map3k1 (cells were cultured in 1 μg/ml Dox for 24 hr prior to analysis). Scale bar: 5 μm; insets: 1 μm. **C)** Immunofluorescence staining of microtubule organization before, during, and 20 min after cold-induced depolymerization of microtubules in RPE1 GFP-Map3k1 cells. Scale bar: 10 μm. See also Fig. 2F. **D)** Immunofluorescence analysis of ciliation (assessed by ARL13B staining) in RPE1 cells inducible expressing GFP-Map3k1 or kinase-dead (KD) GFP-Map3k1. Ciliation was analyzed following 8 hr treatment with or without 1 μg/ml doxycycline (Dox) followed by 48 hr serum starvation in the continued presence/absence of Dox. Bars show means from $N$ = 3 replicate experiments. *** denotes $P$ < 0.001. **E)** Representative U-ExM image of RPE1 cells over-expressing GFP-Map3k1-KD. Cells were stained for tubulin and GFP. Scale bar: 1 μm. **F)** Analysis of cell cycle stage distribution by propidium iodide staining and flow cytometry is shown for the indicated cell lines grown in the presence or absence of doxycycline (Dox) for 24 hr. Asterisks denote significant (**, P < 0.01; ***, P < 0.001) differences in mean of $N$ = 3 replicate experiments. NS = not significant.

### Figure S3

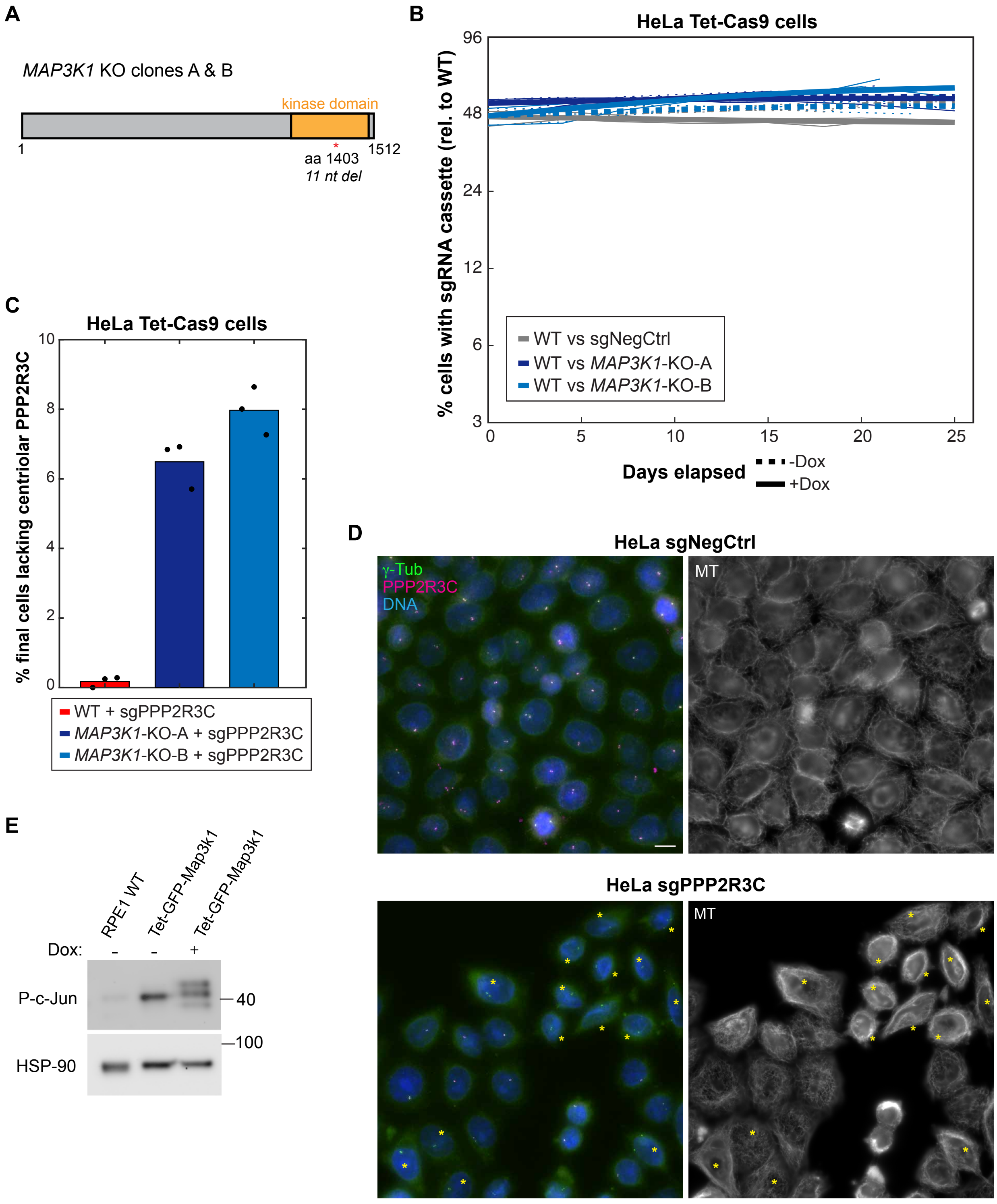

A
MAP3K1 KO clones A & B
kinase domain
1
1512
aa 1403
11 nt del
B
HeLa Tet-Cas9 cells
% cells with sgRNA cassette (rel. to WT)
96
48
24
12
6
3
0
5
10
15
20
25
WT vs sgNegCtrl
WT vs MAP3K1-KO-A
WT vs MAP3K1-KO-B
Days elapsed
-Dox
+Dox
C
HeLa Tet-Cas9 cells
% final cells lacking centriolar PPP2R3C
10
8
6
4
2
0
WT + sgPPP2R3C
MAP3K1-KO-A + sgPPP2R3C
MAP3K1-KO-B + sgPPP2R3C
D
HeLa sgNegCtrl
γ-Tub
PPP2R3C
DNA
MT
HeLa sgPPP2R3C
MT
E
RPE1 WT
Tet-GFP-Map3k1
Tet-GFP-Map3k1
Dox:
-
-
+
P-c-Jun
40
100
HSP-90

**Figure S3. Characterization of *PPP2R3C* and *MAP3K1* knockout HeLa cells**

**A)** Illustration of mutations validated by Sanger sequencing of RPE1 *PPP2R3C* and *FOP* KO cell lines generated by CRISPR. **B)** Flow cytometry growth competitions to assess growth rates (relative to wildtype parental HeLa Tet-Cas9 cells) of HeLa cells transduced with a non-targeting control sgRNA (sgNegCtrl) or HeLa *MAP3K1* KO clones. **C)** Immunofluorescence analysis of PPP2R3C localization at centrioles was assessed for growth competitions shown in Fig. 3D at the final timepoints (note: percentages shown indication the fraction of all cells that lack PPP2R3C signal at centrioles and that at these times the fraction of mCherry-positive cells is 1-10%). Bars show means from $N = 3$ independent experiments. **D)** Immunofluorescence analysis of microtubules, γ-tubulin (γ-tub), and PPP2R3C in HeLa Tet-Cas9 cells transduced with a non-targeting control sgRNA (sgNegCtrl) or an sgRNA cassette targeting *PPP2R3C*. Cells were analyzed after 8 days of Cas9 induction. Asterisks denote cells apparent *PPP2R3C* KO cells (cells lacking centriolar PPP2R3C staining). Scale bar: 10 μm. **E**) Immunoblot analysis of phosphorylated c-Jun (P-c-Jun) and total c-Jun levels in whole cell lysates prepared from the RPE1 wildtype or Tet-GFP-Map3k1 cells grown for 1 hr in the presence or absence of 2 μg/ml Nocodazole (Noc) and for the preceding 24 hrs in the presence or absence of 1 μg/ml Dox. HSP90 was used as a loading control. One of $N = 3$ replicate experiments.

**Figure S4**

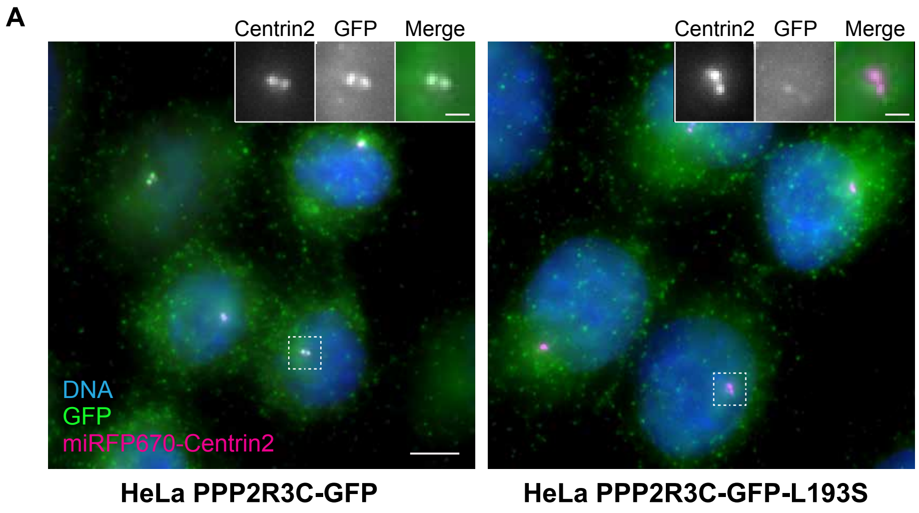
A
Centrin2
GFP
Merge
DNA
GFP
miRFP670-Centrin2
HeLa PPP2R3C-GFP
Centrin2
GFP
Merge
HeLa PPP2R3C-GFP-L193S

**Figure S4. A pathogenic PPP2R3C mutant is defective in centriolar localization**

**A)** Immunofluorescence analysis of PPP2R3C-GFP and miRFP670-Centrin2 in HeLa cell lines stably co-expressing miRFP-670-Centrin and wildtype or L193S forms of PPP2R3C-GFP. Scale bar: 5 μm; insets: 1 μm.
